# Characterization of the growth and morphology of a BSL-2 *Coccidioides posadasii* strain that persists in the parasitic life cycle at ambient CO_2_

**DOI:** 10.1101/2022.03.29.486294

**Authors:** Javier A. Garcia, Kiem Vu, George R. Thompson, Angie Gelli

## Abstract

*Coccidioides* is a dimorphic fungus responsible for Valley Fever and is the cause of severe morbidity and mortality in the infected population. Although there is some insight into the genes, pathways, and growth media involved in the parasitic to saprophytic growth transition, the exact determinants that govern this switch are largely unknown. In this work, we examined the growth and morphology of a novel *Coccidioides posadasii* strain (*C. posadasii* S/E) that efficiently produces spherules and endospores and persists in the parasitic life cycle at ambient CO_2_. We demonstrated that *C. posadasii* S/E remains virulent in an insect infection model. Surprisingly, under spherule-inducing conditions, *C. posadasii* S/E culture was found to be completely hyphal. Differential interference contrast (DIC) and transmission electron microscopy (TEM) revealed unexpected cellular changes in this strain including cell wall remodeling and formation of septal pores with Woronin bodies. Our study suggests that the *C. posadasii* S/E strain is a useful BSL-2 model for studying the molecular factors that are active during the parasitic life cycle and the mechanisms underlying the parasitic to saprophytic growth transition – a morphological switch that can impact the pathogenicity of the organism in the host.

## Introduction

Dimorphic fungi are major causes of fungal disease particularly due to their adapted life cycles in the environment and the hosts. *Coccidioides* is a major cause of pulmonary disease in the arid Southwestern United States where the disease manifests in patients’ lung and can disseminate to the skin, bones, and central nervous system ^1,2^. Infection begins when arthroconidia are inhaled through exposure sites in endemic areas. Although *Coccidioides*’ parasitic life cycle has been largely characterized by the spherule/endospore (SE) phase in the host, much less is known about its hyphal morphologies *in vivo*.

*Coccidioides posadasii* and *Coccidioides immitus* are two known species responsible for coccidioidomycosis, an opportunistic fungal infection that causes significant morbidity and mortality in endemic areas ^1,3^. Although located in the Southwestern part of the United States, the exact distribution of *Coccidioides* in the soil is difficult to determine even in endemic areas ^4^. Furthermore, the factors that favor growth in the environment and within the host remain largely unknown. Increasing concentrations of greenhouse gases and rising temperatures promote *Coccidioides*’ thermotolerance and survival in the environment -traits that can help the organism adapt to growth in the human host ^3-5^.

The factors that regulate *Coccidioides* dimorphic life cycle have been implicated with RNA-seq and proteomics data^6^. Previous studies used global transcriptomics to examine differentially expressed genes in the saprophytic (hyphal) and parasitic (spherule/endospore) life cycles^7^. However, the exact mechanisms underlying the morphological switch remain elusive. Here, we examine the growth dynamics, morphology, and ultrastructures of a novel strain of *Coccidioides posadasii* (*C. posadasii* S/E) that persists in the parasitic life cycle in Converse media at ambient CO_2_ but switches to saprophytic growth (hyphal morphology) in media that mimics an *in vivo*-like environment, a phenomenon not expected based on previous reports^8^. Our characterization of the *C. posadasii* S/E strain that maintains its spherule/endosphore life cycle under routine laboratory condition provides the *Coccidioides* research community with a model organism for studying the molecular factors that are active during the parasitic life cycle and the mechanisms underlying the parasitic to saprophytic growth transition – an important morphological change that has the potential to impact the pathogenicity of the organism.

## Materials and Methods

### Strains and growth conditions

A Silveira strain of *Coccidioides posadasii* was maintained in the spherule/endospore phase (parasitic phase) and approved by Biological Use Authorization for Biosafety Level 2 use. *C. posadasii* spherules stored at -80°C in 15% glycerol were inoculated into 150 mL of chemically-defined Converse media. Converse media, prepared as previously described, contained potassium phosphate monobasic, zinc sulfate, calcium chloride dihydrate, sodium chloride, sodium bicarbonate, magnesium sulfate heptahydrate, ammonium acetate, dextrose, Tamol, and potassium phosphate dibasic trihydrate, filter-sterilized with 0.22 µM Stericup (Millipore) and stored at room temperature ^9,10^. Cultures were grown in vented flasks, incubated at 37°C at ambient CO_2_/O_2_ and shaken at 160 rpm for up to 7 days as previously described^8^. The initial culture grown in Converse medium was sub-cultured into the respective test media by transferring 100 uL aliquots of the initial culture to 150 mL of the respective test media. Roswell Park Memorial Institute-1640 (RPMI-1640) media (10.2 g/L) was supplemented with 10% heat-inactivated fetal bovine serum (FBS) (Life Technologies) and 0.08% Tamol® and filter-sterilized with 0.22 µM Stericup (Millipore) to create RPMI-sph media and stored at 4°C for the continuous production of spherules and co-cultures with mammalian cell lines ^8,11^. RPMI-tamol media contains only RPMI-1640 (10.2 g/L) and 0.08% Tamol® with no added FBS.

### *Galleria mellonella* killing assay

The *G. mellonella* killing assays were carried out as previously described but modified for *Coccidioides* ^12,13^. Briefly, *G. mellonella* wax moth caterpillars (or larvae) (Vanderhorst, Inc., St. Marys, Ohio) that were in the final instar stage were housed in the dark and used within 14 days from the day of shipment. Fourteen caterpillars of the desired weight (∼245 mg +/-25mg) were used in all assays. The inocula were prepared as follows: *C. posadasii* S/E was grown for approximately 144 h in 100 mL of Converse media. The culture was centrifuged at 2000 rpm for 2 mins and washed with 1X PBS three times. Cultures were visually inspected by light microscopy to ensure no contamination was present. Spherules were resuspended in 1X PBS, lightly mixed to break up cluster in order to accurately determine inoculum size and measured with a hemocytometer (Bright-Line™ Hemacytometer, Sigma-Aldrich). Final concentrations of 1.25×10^3^ and 1.25×10^5^ cells/μL was achieved for injection. Before injection, the Hamilton syringe was cleaned sequentially with 10% bleach, 70% ethanol, and 1X PBS. Larvae were injected with 8 μL aliquots of the respective inoculum (10^4^ and 10^6^ total spherules per larvae) in the last left proleg (hemocoel) with the syringe. Injection area was cleaned with an ethanol swab and ampicillin (20 mg of ampicillin/kg of body weight) was co-administered to prevent contamination from native bacteria. Control larvae were injected with 8 μL of 1X PBS or heat-killed spherules of the same concentration. Spherules were inactivated by modifying a previous described procedure ^14^. Briefly, aliquots were boiled in 100ºC water bath for 30 min and cell death was verified using 0.4% trypan blue solution (Millipore Sigma). Larvae were incubated in 37ºC post-injection and monitored daily for survival. Graphs and statistical analysis were done with GraphPad Prism 5.0 statistical software (GraphPad, San Diego, CA). Differences in survival (log rank and Wilcoxon tests) were analyzed by the Kaplan-Meier method. P values of <0.05 were considered significant^11^. Leica DIC DMRE imaging.

Cultures were grown in different media for 7 days in the 37°C shaking incubator, washed with 1X PBS and fixed for one hour in 4% paraformaldehyde. Spherules were rinsed off with 1X PBS and mounted on a coverslip. Differential Interference Contrast (DIC) images were taken on a Leica DMRE microscope running MetaMorph v7.1. Images were processed on ImageJ. DIC images taken were also used to quantitatively compare spherule diameter in Converse and RPMI-tamol media. For the quantification of spherule diameter, ImageJ was calibrated with a micrometer (Reichert-Jung, Leica) and values were recorded for each media condition. Sample size for each group was 30 spherules. Graphs and statistical analysis were done with GraphPad Prism statistical software using an unpaired t-test with Welch’s correction (GraphPad, San Diego, CA). P values of <0.05 were considered significant.

### Enzymatic release of extracellular proteins from S/E phase *Coccidioides*

Spherules were harvested after 144 -168 h and washed with phosphate buffered saline (PBS). Spherule surface proteins were released in a 15-mL tube in triplicate by treating intact spherules with 3 mL of 0.25% trypsin/EDTA solution (Corning) for 4 h or 14 h at 37ºC and room temperature, respectively, on a rotating mixer. Following the trypsin treatment, the spherules were examined by compound light microscopy to ensure that they remained intact. Peptides resulting from the trypsin treatment were concentrated using a 3 kDa cutoff filter (Amicon) and separated on an SDS-PAGE gel and visualized with Coomassie blue stain. For comprehensive trypsin digests, complete and concentrated samples were sequenced by LC-MS/MS.

### Mass Spectrometry Sequencing

Supernatant from trypsinized cells and proteins bands of interest were excised from polyacrylamide gels and submitted to the UC Davis Proteomic Core. Scaffold (Proteome Software Inc., Portland, OR) was used to validate MS/MS based peptide and protein identifications.

### Transmission electron microscopy

Cultures were grown in Converse, RPMI-tamol, and RPMI-sph media for 7 days in the 37°C shaking incubator, washed three times with 1X PBS and fixed in modified Karnovsky’s fixative (2.5% glutaraldehyde, 2% paraformaldehyde in 0.1M sodium phosphate buffer) overnight. Samples were submitted to the Biological Electron Microscopy Facility at University of California, Davis for TEM sample preparation. Briefly, cells fixed overnight were centrifuged (2000 rpm for 3 min) to remove the fixative and rinsed in 0.1 M sodium phosphate buffer. Cells were then fixed in 1% osmium tetroxide/1.5% potassium ferrocyanide for 1 h and rinsed with cold water. Cells were dehydrated with 30%, 50%, 70%, 95%, and 100% (3 times) ethanol for 10 mins each (7 total). Resin (Dodecenyl Succinic Anhydride, Araldite 6005, Epon 812, Dibutyl Phthalate, Benzyldimethylamine) was added and allowed to infiltrate cells overnight at room temperature. Fresh resin was added the next day and set to polymerize at 70°C overnight. Polymerized blocks were sectioned on a Leica EM UC6 ultramicrotome at approximately 100 nm and collected on a copper grid. Grids were dried in 60°C oven for 30 min and stained with 4% aqueous uranyl acetate and 0.4% lead citrate in 0.1N NaOH. Sections were imaged with a TEM microscope (FEI Talos L120C TEM 80kv) and images processed on ImageJ.

### *In vitro* growth kinetics

Fresh 150 mL of test media (Converse, RPMI-tamol, RPMI-sph) were inoculated with 100 μL of a 7-day initial S/E culture grown in Converse media. Cultures were visually inspected using light microscopy to ensure no contamination was present. Growth rates were determined by measuring OD_600_ and blanking the spectrophotometer with the respective media. Cultures were mixed prior to sampling to ensure settled cultures were homogenous. Measurements were taken in triplicate every 24 h for a period of 17 days starting at time zero. Light microscopy images were taken at the end of the 17 day incubation period to show a representation of the culture’s morphology

## Results

### *C. posadasii* S/E spherules are virulent *in vivo*

We investigated the virulence of *C. posadasii* S/E spherules in the *Galleria mellonella* insect model of infection. Survival of larvae infected with 10^4^ live spherules was not different from larvae inoculated with PBS or with 10^4^ heat-killed spherules (*P* = 0.7786) (Figure 1A). Since the size of the inoculum can have varying effects on the larvae, we next challenged *G. mellonella* larvae with 10^6^ spherules. At this inoculum size, we observed a statistically significant reduction in percent survival of larvae infected with live spherules compared to equal amounts of heat-killed spherules (*P* = <0.0001) or PBS-only control (P = <0.0001) (Figure 1B). Larvae challenged with 10^6^ spherules succumbed to *C. posadasii* S/E within 4 days of infection (Figure 1B). On the other hand, 95% of larvae challenged with spherules that had been heat-killed were alive on day 4 and by the 8th day post-infection, 30% of animals survived (Figure 1B). Improvement in the survival of larvae inoculated with heat-killed *C. posadasii* S/E spherules suggested that the mechanism(s) of virulence was partly proteinaceous-based. Similarity in surface antigens of *Coccidioides* identified in human chest tissue and *C. posadasii* S/E strain.

**Figure 1.**
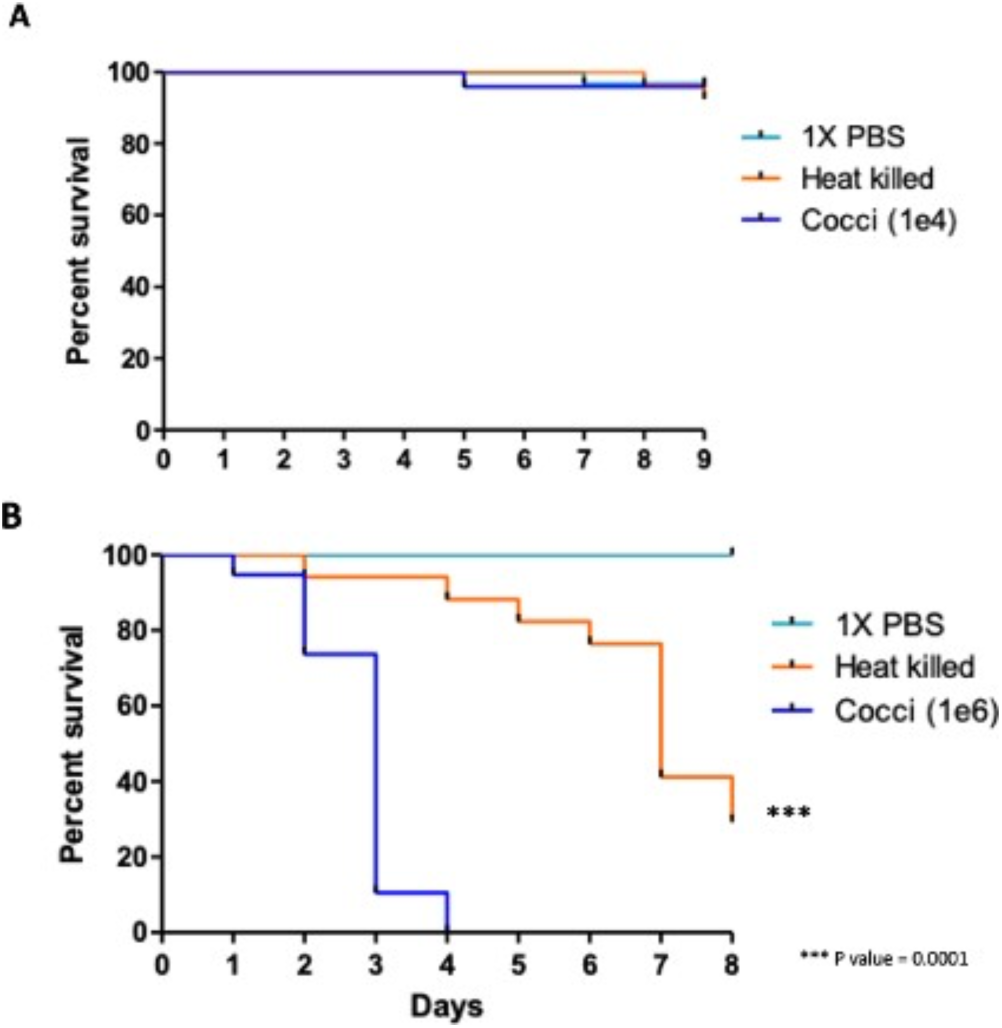
The *C. posadasii* S/E strain is virulent inan insect model of infection. Larva of *Galleria mellonella* inoculated with 10^4^ spherules were notaffected (**A**), whereas inoculation of 10^6^ spherules resulted in 100% death of larvae within4 days (**B**). Heat-inactivation of the *C. posadasii* strain prior to inoculation improved survival significantly suggested that the mechanism of virulence is proteinaceous-based. PBS controls show no difference in survival. Statistical analysis was performed using GraphPad Prism software by estimating differences in survival (log rank andWilcoxon tests) using the Kaplan-Meier method. Shown is one of three replicates.

To further assess the virulence of *C. posadasii* S/E, we evaluated the surface antigens of the spherules using a *Coccidioides*-specific monoclonal antibody (C-mAb) raised against a surface antigen. We found that C-mAb identified spherules in human chest tissue from a confirmed case of coccidioidomycosis (Figure 2, top panel). The C-mAb also stained the surface of *C. posadasii* S/E spherules that had been cultured and fixed (Figure 2, bottom panel).

**Figure 2.**
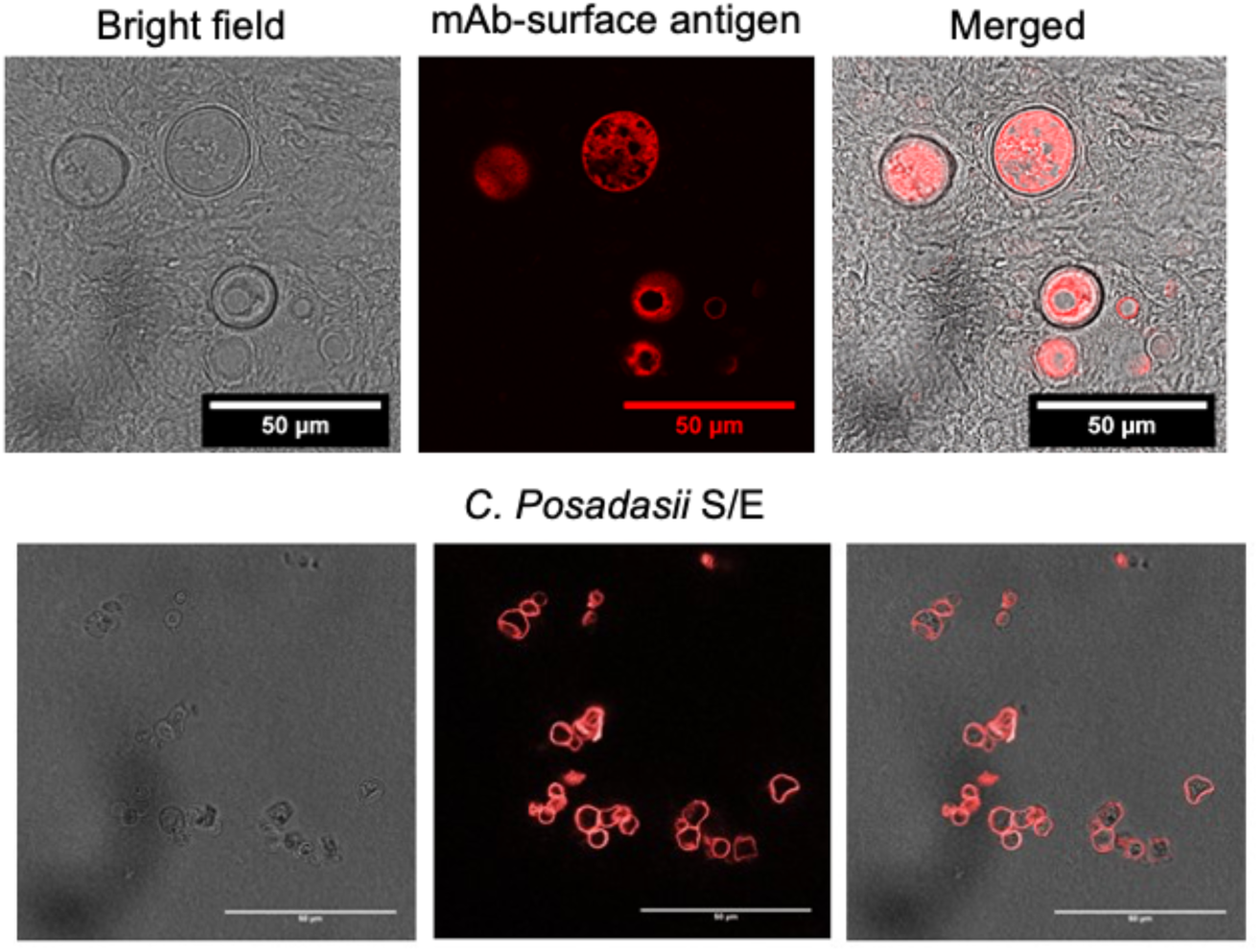
Surface antigen of *C. posadasii* S/E strain similar to *Coccidioides* identified in human chest tissue. Top panel: IHC staining with a *Coccidioides* surface antigen mAb detects spherules in human chest tissue with confirmed coccidioidomycosis. *Coccidioides* spherules were identified by brightfield imaging 10-30 µm. IHC staining using the *Coccidioides* mAb primary antibody and Alexa Fluor® 568 secondary antibody showed specific fluorescent signal localized on the surface of spherules (100x magnification). Brightfield and fluorescence images were merged and showed co-localization of spherules and fluorescent signal.Bottom panel: the same primary mAb used above stainedthe surface of *C. posadasii* S/E stain fixed with thimerosal. *Coccidioides* mAb primary antibody and Alexa Fluor® 568 secondary antibody showed specific fluorescent signal localized on the surface of spherules.

The *Coccidioides*-specific mAb did not cross react with a strain of *Cryptococcus neoformans*, healthy human lung/chest tissue or diseased lung tissue (large cell carcinoma T2A N0 G3 tumors or squamous cell carcinoma) (data not shown). These results suggested that surface antigens of *C. posadasii* S/E spherules are likely similar to those found on wild-type *Coccidioides* spherules, consistent with the virulence of C. *posadasii* S/E in the *G. mellonella* model of infection.

We further examined the surface antigens of *C. posadasii* S/E spherules by identifying surface-exposed proteins. We isolated 4 highly abundant extracellular proteins by utilizing trypsin shaving of intact spherules from the cultured *C. posadasii* S/E strain and mass spectrometry for protein identification. Copper-zinc superoxide dismutase (SOD), DOMON-like type 9 carbohydrate binding module, aspartyl protease and mannosyl-oligosaccharide alpha-1,2-mannosidase precursor were identified as surface proteins of *C. posadasii* S/E spherules (Table 1). These same proteins were identified as immunoreactive antigens in other *Coccidioides* strains from previous studies ^15-19^.

**Table 1.** Immunoreactive proteins on the surface of *C. posadasii* S/E spherules. Surface proteins were isolated by Trypsin shaving of intact spherules, separated by SDS-PAGE electrophoresis and excised polypeptide band was identified by mass spectrometry.

| Immunoreactive proteins identified |  |  |  |  |
| --- | --- | --- | --- | --- |
| Protein/Predicted function | Accession number | Alternate ID | Molecular weight (kDa) | Previous reports |
| Copper, zinc superoxide dismutase (SOD) | C5P5N2_COCP7 | CPC735_033580 | 26 | [15] |
| DOMON-like type 9 carbohydrate binding module | C5PIH6_COCP7 | CPC735_056990 | 26 | [16] |
| Aspartyl protease | C5P9L1_COCP7 | CPC735_005950 | 44 | [17, 18] |
| Mannosyl-oligosaccharide alpha-1,2-mannosidase precursor | C5PAF0_COCP7 | CPC735_008870 | 57 | [19] |

### *C. posadasii* S/E reverts to mycelial form in the spherule-producing media, RPMI-sph

As expected, *C. posadasii* S/E produced spherules and endospores continuously and rapidly in Converse media at ambient CO_2_/O_2_ without the need to initiate the culture from arthroconidia (Figure 3A, top panel). Prior work found that wild-type *Coccidioides* strains continuously produce spherules in RPMI-sph media, which provides conditions similar to that found *in vivo*^6,9^. Therefore we wanted to ask whether *C. posadasii* S/E also behaves like wild-type strains in RPMI-sph. Surprisingly, we found that *C. posadasii* S/E reverted to a nearly 100% hyphal form without any observable arthroconidia after 7 days of culture in RPMI-sph (Figure 3A, bottom panel).

**Figure 3.**
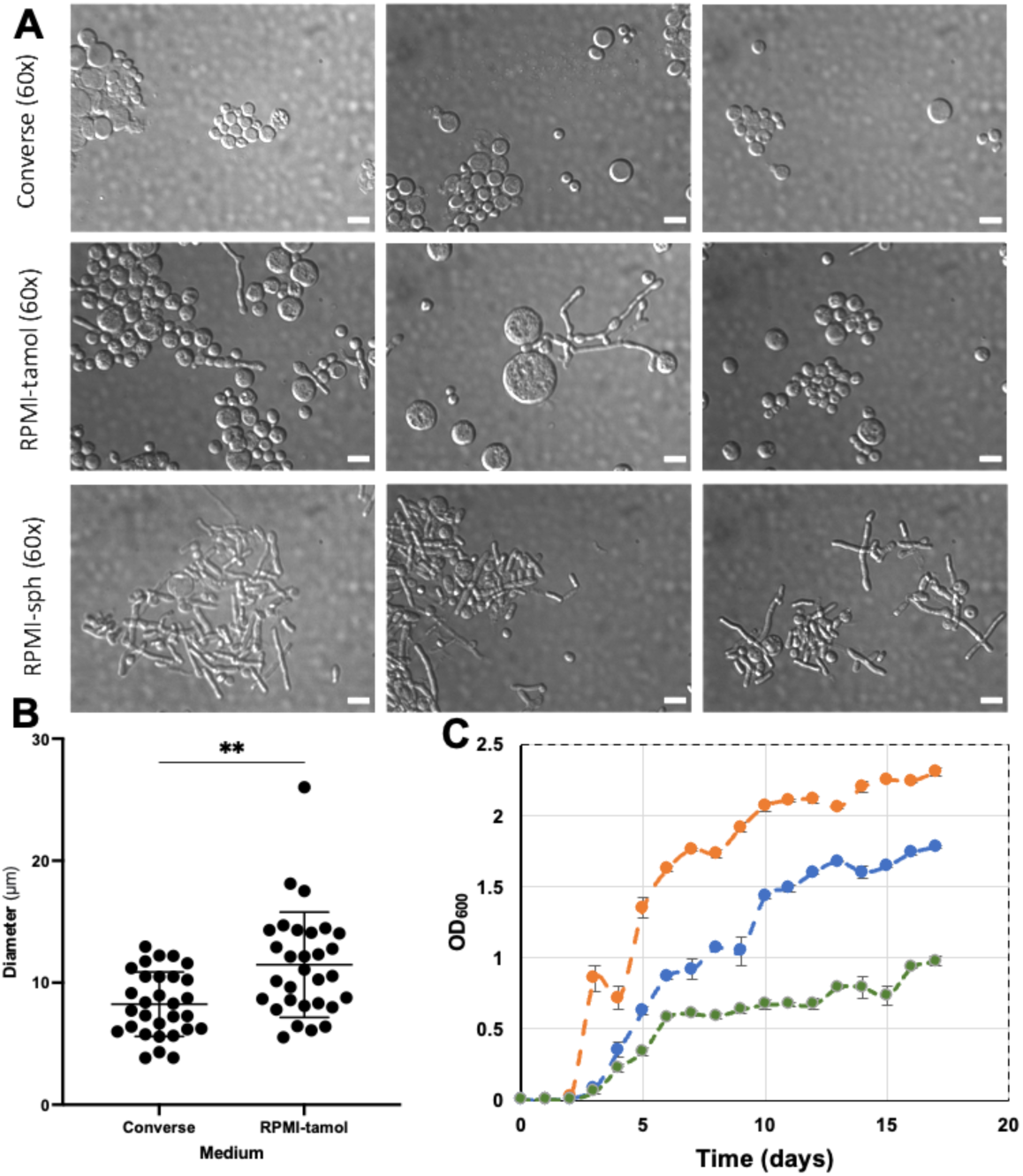
Differential Inference Contrast (DIC) microscope images show *C. posadasii S/E morphology* and size indifferent types of media. **A**: Spherule and endospore production is optimal in Converse media, a mixture of spherule/endospores and hyphal morphologies in RPMI-tamol media, and complete hyphal morphology in RPMI-sph media. Images taken in triplicate on 60x magnificationwith Leica DMRE microscope. Scalebar represents 10 µm. **B**: Spherules grown in RMPI-tamol are more varied and larger on average than spherules grown in Converse media. Graphs and statistical analysis were done using GraphPad Prism statistical software using an unpaired t-test with Welch’s correction (*P* = 0.001), n=30. **C**: The growth rates of the strain in 3 different mediums: Orange – RPMI-sph; Blue – converse media; Green -RPMI-tamol.

To identify which media component may have contributed to the transition from parasitic (spherule/endospore) to saprophytic (environmental form) growth, we removed FBS from RPMI-sph to create RPMI-tamol. A mixture of both mature spherules and mycelia was observed when *C. posadasii* S/E was grown in RPMI-tamol (Figure 3A, middle panel). We also noted that *C. posadasii* S/E spherules from RPMI-tamol culture were larger and consistently more heterogenous in size compared to those grown in Converse media (Figure 3B). Therefore, we asked whether there was a difference in the growth rate of *C. posadasii* S/E in the 3 different media (Figure 3C). Robust growth was observed in RPMI-sph relative to growth in Converse or RPMI-tamol. Among the three test media, *C. posadasii* S/E grew the slowest in RPMI-tamol (Figure 3C). Ultrastructural features of *C. posadasii* S/E when cultured in different media.

The observed morphological changes of *C. posadasii* S/E in the different media prompted us to examine the ultrastructural features of the strain. Transmission electron microscopy (TEM) revealed enhanced endospore production in both Converse (Figure 4A) and RPMI-tamol media (Figure 4B) and very little spore production in RPMI-sph (Figure 4C). Mycelial cultures grown in RPMI-sph were largely devoid of endospores despite having a growth advantage over cultures grown in either Converse or RPMI-tamol. We also observed changes in the outer cell wall glycoprotein layer. TEM micrographs revealed that spherules grown in both Converse and RPMI-tamol had a loose outer cell wall layer that appeared to shed (Figure 4A,4B). However, micrographs of mycelia grown in RPMI-sph show a consistently smoother outer cell wall with minimal shedding (Figure 4C).

**Figure 4.**
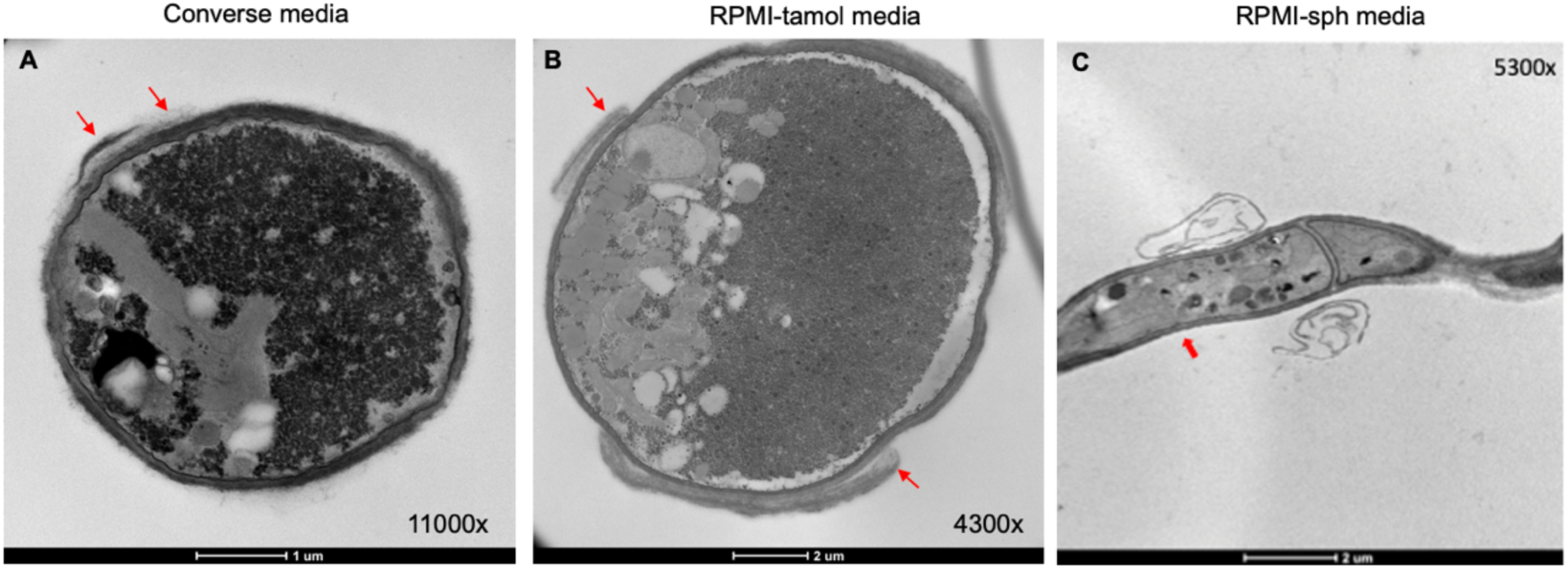
Ultrastructural features of *C. posadasii* S/E strain. TEM revealed a sloughing off of the outer cell wall glycoprotein layer of sperules (red arrows) grown in Converse or RPMI-tamol media that was not observed in the hyphal form (red arrow). Hyphae also appeared to lack endospores in comparison to the spherule. TEM images of *C. posadasii* S/E strain grown in Converse, RPMI-tamol or RPMI-sph media were taken after 168 h growth *in vitro*.

The TEM studies also revealed that spherules grown in RPMI-tamol had ruffled, uneven cell walls. The most striking observation was the frequency of irregularly shaped spherules found in RPMI-tamol compared to the mostly round spherules observed in Converse media (Figure 5). Many spherules found in RPMI-tamol were elongated into an elliptical shape and some appeared to be connected as pseudohyphae (Figure 5).

**Figure 5.**
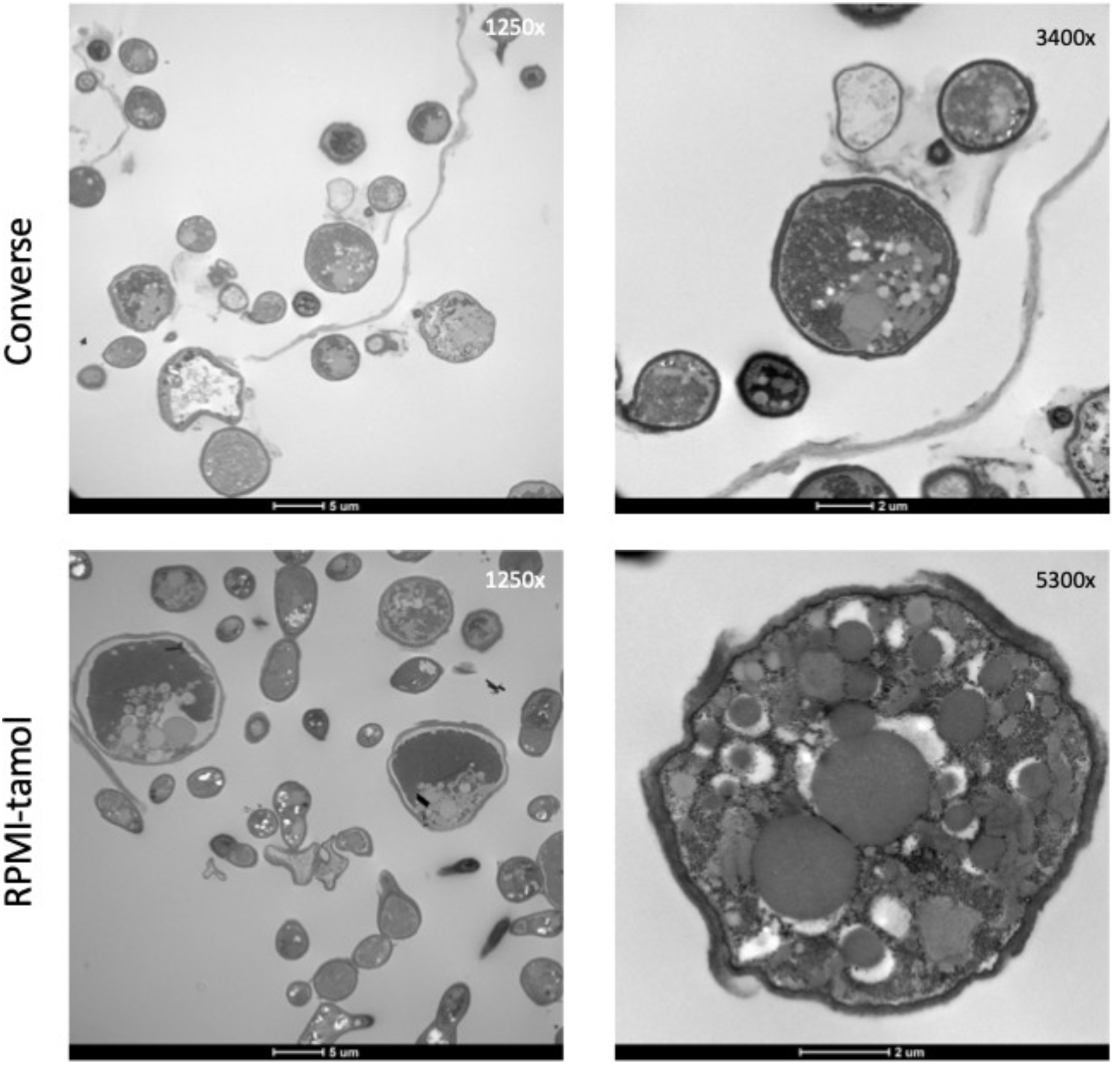
TEM images of *C. posadasii* S/E strain following 168 h growth in RPMI-tamol or RPMI-sph media. The shape of spherules grown in RPMI-tamol media are highly irregular compared to spherules grown in Converse media.

A closer look at the hyphae of *C. posadasii* SE grown in either RPMI-tamol or RPMI-sph revealed a lack of endospores, as expected, and organelles such as mitochondria and nuclei could be identifed (Figure 6). The outer cell wall, inner wall and plasma membrane were also clearly visible. Invaginations at the septum appeared to create a structure resembling a septal pore. The similarly sized organelles near the septal pores resembled Woronin bodies; these are small organelles used by filamentous ascomycetes to plug septal pores after injury (Figure 6)^17^.

**Figure 6.**
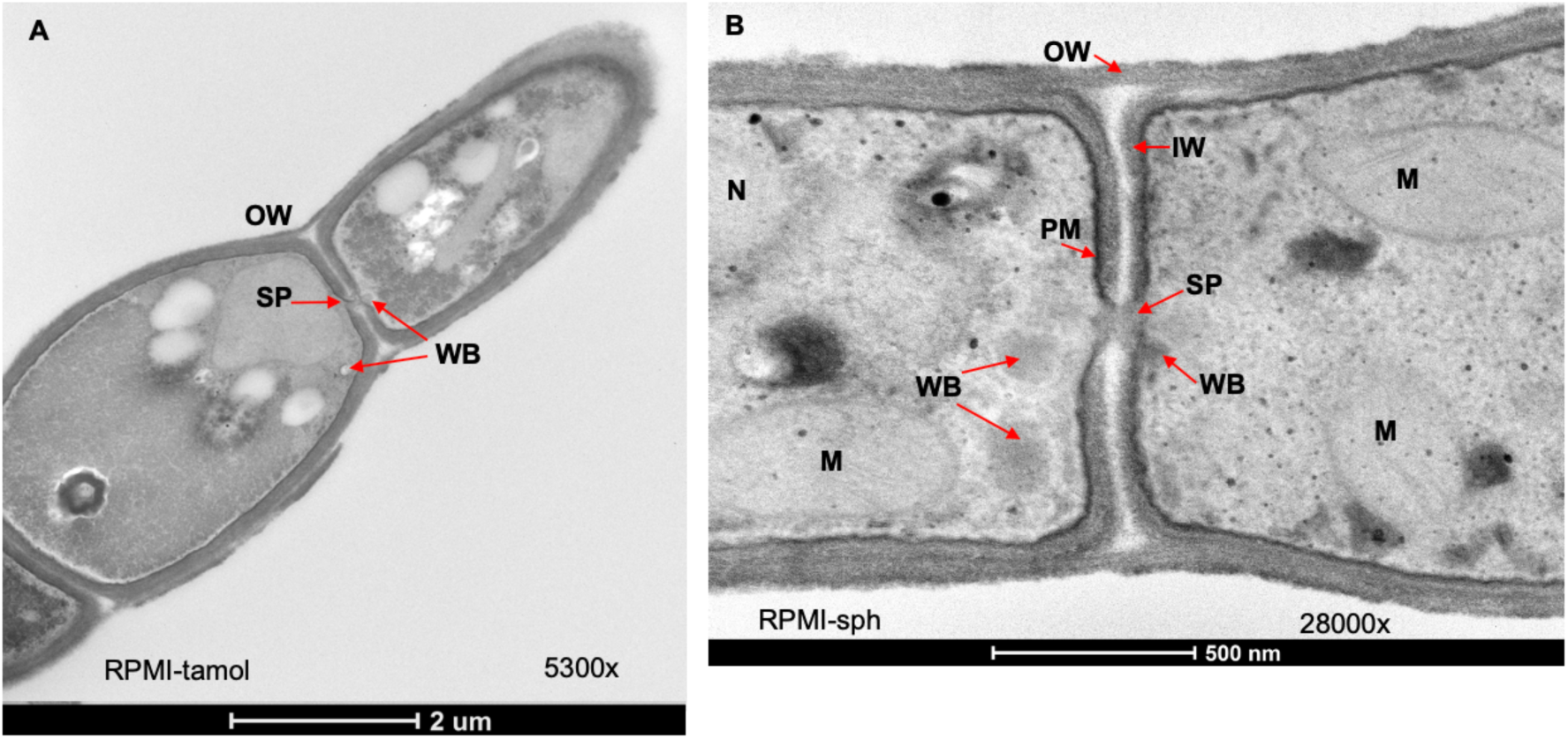
Hallmarks of mycelia observed in the mycelial form of *C. posadasii* S/E strain. TEM images of *C. posadasii* S/E strain following 168 h growth in RPMI-tamol or RPMI-sph media. SP: sepatal pore; WB: Woronin bodies; OW: outer wall; IW: inner wall; PM: plasma membrane; M: mitochondria; N: nucleus.

## Discussion

The aim of this study was to characterize the novel *C. posadasii* S/E strain and to establish whether it could serve as a useful BSL-2 model of *Coccidioides*. Whereas wild-type strains of *Coccidioides* require 5-10% CO_2_ for spherule induction in RPMI-sph, *C. posadasii* S/E could readily produce pure cultures containing spherules and endospores in Converse media at ambient CO_2_/O_2_ with no observable hyphae. Additionally, arthroconidia is not required to initiate *C. posadasii* S/E cultures, which can readily be started from a spherule glycerol stock. *C. posadasii* S/E has been approved by Biological Use Authorization for Biosafety Level 2 use because the strain strongly persists in the parasitic life cycle under routine laboratory conditions and there is low risk associated with unwanted sporulation. The availability of *C. posadasii* S/E for the research community would greatly facilitate future work on this under-studied human fungal pathogen because of the ease of producing large quantity of spherules for research and the ability to carry out studies under BSL-2 conditions.

Previous work utilized RPMI-containing media for the co-cultivation of both spherules and mammalian cell lines as well as the conversion of arthroconidia to spherules. A large body of literature has focused on the media composition that promotes the transition from saprophytic to parasitic growth since this morphological switch is important for the virulence of *Coccidioides* in the human host^7,9,18-27^. Significant difference in spherule density and size among various *Coccidioides* strains grown in RPMI-sph media were reported^6^. In contrast, *C. posadasii* S/E culture was found to be completely hyphal in RPMI-sph, and we did not observe new spherule formation even after 17 days of incubation. Rare spherules could be found in the RPMI-sph culture, but they likely originated from the initial inoculation used to start the culture.

Unlike the observed uniform mycelial culture of *C. posadasii* S/E in RPMI-sph, Mead et al. and Petkus et al. found both mycelia and spherules in RPMI-sph cultures of several *Coccidioides* strains^8,10^. Mycelia from cultures of *C. posadasii* C735, *C. posadasii* C735 Δcts2/Δard1/Δcts3, and *C. immitus* RMSCC 2006 could be detected at a later time (days 7-10); however, there was still significant spherule formation during this incubation period ^8^. In contrast to the uniform mycelial morphology of *C. posadasii* S/E in RPMI-sph, *C. posadasii* S/E culture grown in RPMI-tamol had a mixture of both spherules and mycelia, and the spherules were significantly larger than those obtained from cultures grown in Converse media after 7 days of incubation. This result echos the finding from the Mead et al. study in which spherules from wild-type *Coccidioides* strains were found to be larger in RPMI-tamol compared to those grown in Converse media. Together, our work and prior studies suggest that *C. posadasii* S/E behaves like a wild-type strain in RPMI-tamol^6^ but in RPMI-sph, *C. posadasii* S/E adopts a uniform hyphal morphology.

FBS is known to stimulate optimal hyphal growth of *C. albicans* at 37ºC by downregulating a major regulator of hyphal morphogenesis, Ras1, suggesting that the Ras1-Cyr-PKA pathway might play a role in regulating the life cycles of dimorphic fungi ^20,21^. Furthermore, the Ras1-Cyr-PKA pathway is involved in activating the transcription factors Cph1 and Efg1, both of which are important for hyphal formation and activation of hyphal molecular switches in *C. albicans* ^22^. To resolve the different regulators involved in the conversion of spherules to hyphae in the presence of FBS, transcriptomic and proteomic studies could potentially clarify the observed differences among different strains of *Coccidioides*.

A body of literature has documented the rare mycelial parasitic morphology of *Coccidioides* in several patients that present mycelial forms in tissues ^23-29^. It is conceivable that the mycelial forms in the host could potentially be due to localized microenvironments created by low CO_2_ conditions, cell-specific stimuli, or varying degrees of immunocompetence of the host ^28,30^. This atypical hyphal form can complicate diagnosis since it could be confused with other filamentous fungal pathogens^31,32^. The *C. posadasii* S/E strain could potentially be used as a model organism for studying atypical *Coccidioides* morphology and developing diagnostic tools.

The growth rates of *C. posadasii* S/E varied significantly in different media. Cultivation of *Coccidioides* typically requires a long incubation period: 3-5 days to grow on plates, 6 weeks to harvest arthroconidia, and another 3-7 days to harvest spherules ^7,8^. *C. posadasii* S/E culture reached an OD_600_ of 0.8553 by day 3 in RPMI-sph, by day 6 in Converse, and by day 15-16 in RPMI-tamol. The different growth rates could potentially be attributed to the additional carbon source and nutrients from fetal bovine serum (FBS). We found that the growth curve of *C. posadasii* S/E culture was atypical and characterized by fast changes in optical density, likely as a result of nutrient exhaustion and nutrient shifting. It has been noted that several filamentous ascomycetes including *Penicillium ochrochloron, Trichoderma harzianium, Aspergillus niger*, and *Aspergillus nidulans* have similar growth curves that do not follow the classical logarithmic growth followed by a stationary phase ^33^. On average, the cultures continued to increase in optical density over time; however, the higher OD could result from media evaporation due to long incubation (17 days) at 37ºC.

The virulence of *C. posadasii* S/E was assessed in the *Galleria mellonella* insect model. The insect model is advantageous as an *in vivo* model of infection because of its cost effectiveness, ease of handling and assessing viability ^12,34,35^. Currently, there are no published studies that use *G. mellonella* as a model of infection for *Coccidioides*, particularly with spherules as the inoculum. *C. posadasii* S/E spherules was found to significantly reduce survival of *Galleria mellonella* larvae over time. We did not see a reduction in survival when larvae were inoculated with 10^4^ spherules, a comparable inoculum size used in our prior studies on the virulence *Cryptococcus neoformans*.^13^ Increasing the number of spherules to 10^6^ resulted in 100% death of larvae, consistent with previous work showing that the spherule/endospore phase is less virulent than arthroconidia in murine models of infection ^36^.

One limitation we encountered was performing the *G. mellonella* studies with only the *C. posadasii* S/E strain. As a BSL-2 strain, we had the ability to perform these experiments under BSL-2 conditions, which greatly facilitated our work. Due to limitations that prevented us from utilizing BSL-3 facilities, we did not have the ability to work with a wild-type laboratory strain of *Coccidioides* ^8^. Additional work is needed to examine the virulence of *C. posadasii* S/E spherules together with another wild-type laboratory strain such as the parent strain *C. posadasii* Silveira, *C. posadasii* C735, *C. immitus* RS, and *C. immitus* 2006 ^8^. Another BSL-2 strain that warrants further comparison is *C. posadasii* Δcts2/Δard1/Δcts3 -a mutant with the *C. posadasii* C735 background that produces sterile spherules incapable of endosporulation in the parasitic phase and is avirulent in mice ^37^. Future work utilizing mycelia from *C. posadasii* S/E grown in RPMI-sph should also be evaluated to determine if pre-conditioning *C. posadasii* S/E in RPMI-sph has a significant effect on its pathogenicity.

As hyphae, *C. posadasii* S/E produced little to no endospores in the culture. Some hyphae appeared to grow as a pseudohyphae in RPMI-sph while others formed septa. We were the first to image and document septal pores in the saprophytic phase of *Coccidioides*. Septal pores allow the movement of organelles from cell to cell. When cells detect damage, Woronin bodies block septal pores to prevent the loss of organelles and other cellular contents essential for cell survival ^38^. We were able to identify Woronin-like bodies near the septal pore, consistent with prior work showing Woronin-like bodies near the septum using electron microscopy ^39^. Our data also lends support to the study by Whiston el al. which found Hex1, a gene responsible for the formation of Woronin bodies, was upregulated in mycelia ^7^. Several studies have shown that Hex1-homolog mutants in other filamentous fungi including *Aspergillus fumigatus, Arthrobotrys oligospora, Metarhizium robertsii*, and *Verticillium dahlia* have reduced virulence in their respective host ^40-43^. Deactivating the Woronin-body stress response to injury can potentially be used as an anti-virulence mechanism to treat infections caused by mycelial forms in the host.

Spherules grown in either Converse or RPMI-tamol media have a constant shedding of the outer wall as previously observed in the wild-type Coccidioides C375 strain, suggesting that *C. posadasii* S/E spherules retain wild-type phenotypes under these growth conditions ^44,45^. However, examination of mycelia formed in RPMI-sph showed that the outer wall remained intact with minimal shedding. Interestingly, Mead et al found that spherules of *C. posadasii* C735 Δcts2/Δard1/Δcts3 had thick and intact cell walls, similar to what we found with mycelia of *C. posadasii* S/E ^44^. The pathway involving cts2/ard1/cst3 is therefore likely downregulated in *C. posadasii* S/E when cultured in RPMI-sph and is crucial for cell wall remodeling.

In summary, we report the characterization of a *Coccidioides posadasii* (*C. posadasii* S/E) strain that can serve as a useful BSL-2 model for studying the complex molecular changes involved in the life cycle transition in response to specific stimuli.

## Author Contributions

For Conceptualization, A.G., J.A.G., K.V., G.R.T.; methodology, A.G., J.A.G., K.V., G.R.T.; formal analysis, J.A.G., K.V., A.G.; resources, G.R.T.; data curation, J.A.G.; K.V.; writing—original draft preparation and editing to final manuscript: J.A.G.; K.V.; A.G.; G.R.T.; All authors have read and agreed to the published version of the manuscript.

## Institutional Review Board Statement

Not applicable.

## Informed Consent Statement

Not applicable.

## Data Availability Statement

Not applicable.

## Acknowledgments

We are grateful to the members of the Gelli Lab for their insightful discussions. We thank the UC Davis Biological Electron Microscopy Facility staff for the processing of samples for TEM. We thank the UC Davis Center for Valley Fever for sharing reagents, equipment and assistance. JA Garcia was supported by the NIH T32 HL007013 Training in Comparative Lung Biology and Medicine.

